# Atomistic molecular simulations of A*β*-Zn conformational ensembles

**DOI:** 10.1101/2023.03.23.534009

**Authors:** Julen Aduriz-Arrizabalaga, Xabier Lopez, David De Sancho

## Abstract

The amyloid-forming A*β* peptide is able to interact with metal cations to form very stable complexes that influence fibril formation and contribute to the onset of Alzheimer’s disease. Multiple structures of peptides derived from A*β* in complex with different metals have been resolved experimentally to provide an atomic-level description of the metal-protein interactions. However, A*β* is intrinsically disordered, and hence more amenable to an ensemble description. Molecular dynamics simulations can now reach the timescales needed to generate ensembles for this type of complexes. However, this requires accurate force fields both for the protein and the protein-metal interactions. Here we use state-of-the-art methods to generate force field parameters for the Zn(II) cations in a set of A*β* complexes and combine them with the Amber99SB^*⋆*^-ILDN optimized force field. Upon comparison of NMR experiments with the simulation results, further optimized with a Bayesian/Maximum entropy approach, we provide an accurate description of the molecular ensembles for most A*β*-metal complexes. We find that the resulting conformational ensembles are more heterogeneous than the NMR models deposited in the Protein Data Bank.

## Introduction

Alzheimer’s disease is the leading cause of senile dementia and has over 55 million cases reported by the World Health Organisation as of 2021.^1^ Although the cause of Alzheimer’s is not completely known, a relation with aggregation and deposition of A*β* in neural tissue is widely accepted as a contributing factor to the onset of the disease. Transition metal ions and oxidative metabolism have been proposed to play fundamental roles in the process of aggregation and deposition of A*β*.^2^ The binding of divalent metals, such as copper, iron, and zinc, with disordered fibrilogenic proteins, such as A*β*, influences the aggregation process of the protein, contributing directly to the severity of the neurodegenerative disease.^3^ It has been reported that both the monomeric and oligomeric forms of A*β* are neurotoxic^4^ and that these cations directly influence toxicity. ^5^ Interestingly, the Zn(II) ion concentration in the brain^6^ of around 150 *μ*M is an order of magnitude higher than the ion concentration in blood. Furthermore, even though the Zn(II) concentrations remain relatively constant through adult life, a significantly elevated concentration has been found on the brains of patients affected by Alzheimer’s disease.^7^ Therefore, the role Zn(II) plays in Alzheimer’s disease has become of great interest.

Despite the relevance of protein-metal interactions to a thorough understanding of A*β*-Zn(II) complexes, their details have remained elusive. This is partly due to A*β* being an intrinsically disordered protein (IDP), which makes an ensemble description preferable to the static structures derived from experimental methods such as X-ray crystallography or Nuclear Magnetic Resonance (NMR). ^8^ Additionally, Zn(II) characterization is highly limited by its physico-chemical properties. Out of its three stable isotopes, only ^67^Zn is NMR active, but due to its low natural abundance and low receptivity, only solid-state, low-temperature NMR studies of zinc compounds are practically achievable. In addition, zinc complexes have no absorbance in the UV-Vis and microwave spectral regions, and its completely filled d^10^ orbitals render it diamagnetic, being, therefore, invisible in EPR spectroscopy.^9^ All these factors combined make experimental studies of A*β*-Zn(II) systems highly challenging. Computer simulations are ideally suited to complement experiments in our understanding of these types of systems.

Modelling complexes of A*β* with transition metals like Zn(II) is however not without its own challenges. First, the chemical flexibility of Zn(II) allows it to adopt different coordination modes with A*β* (see Figure 1). In the experimentally resolved structures of A*β* with Zn(II) complexes, we find the metal tetrahedrally coordinated by glutamic/aspartic acid and/or histidine residues, ^10^ making different combinations possible. Hence, modelers have multiple possibilities to simulate the protein-transition metal interactions. ^11^ The simplest option is through nonbonded models, where the metal ion interacts via electrostatic and Van der Waals interactions. Nonbonded models have successfully been used in the past.^12–15^ However, in the case of Zn(II) nonbonded models often result in the wrong coordination, and the metal may even leave the coordination site.^11^ This problem may be particularly acute for complexes with IDPs, where the metal may be more solvent-exposed than in globular proteins. In order to overcome these limitations, one possibility is to use “dummy models”, which contain virtual atoms with point charges that promote the target coordination.^16,17^ Unfortunately, they have been reported to undergo dissociation in long-time scale simulations.^11^ Alternatively, one can use constraints to force the coordination of metal binding.^10,18,19^ Nonetheless, it has been reported that improbable fluctuations appeared using this approach. ^11^ Alternatively, one can use bonded models, where chemical bonds are defined between the protein and the metal, which keep the coordination stable throughout the whole simulation time. Bonded models have been used extensively for zinc-containing systems,^2,20–22^ including A*β*-metal complexes.^3,23–26^

**Figure 1:**
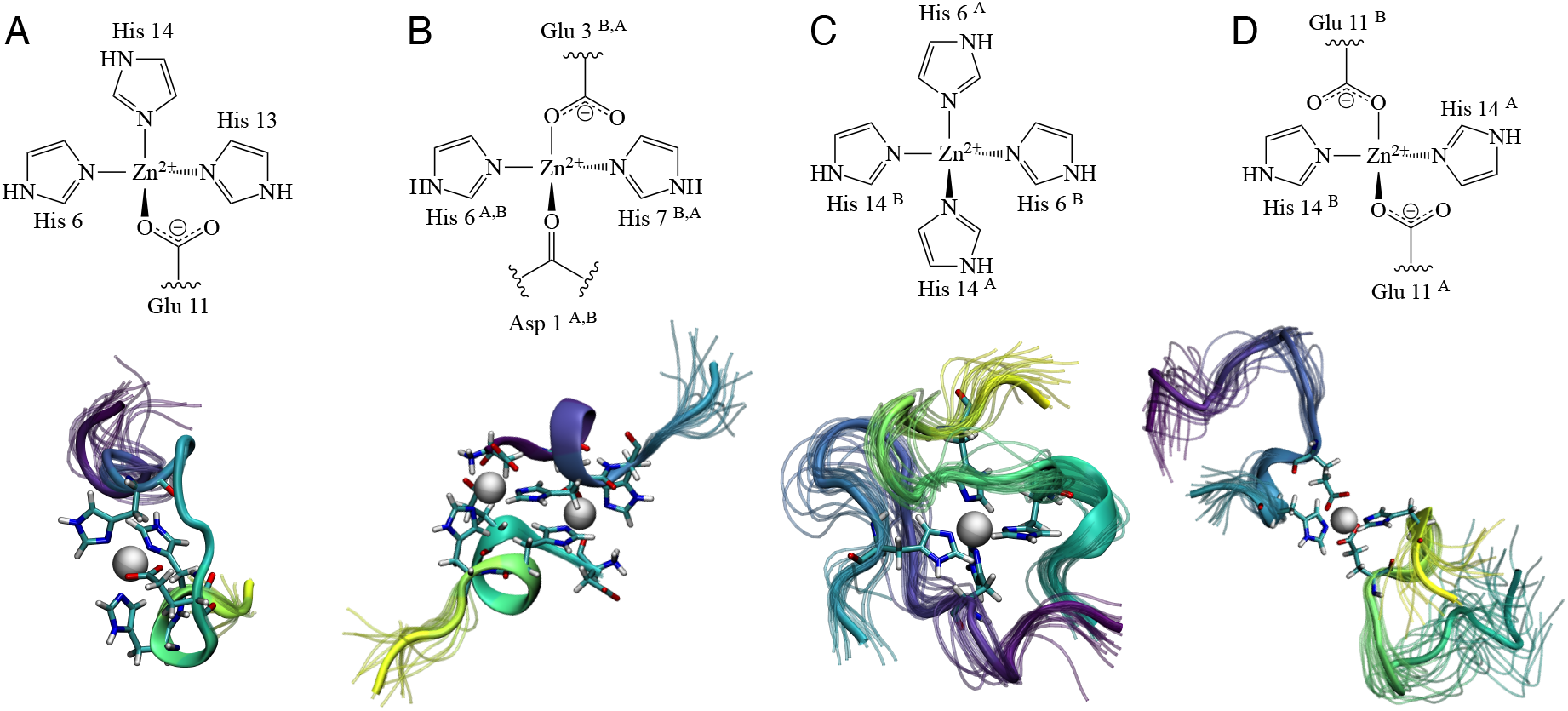
Cartoon representation of the metal centers (top) and cartoon representation of A*β*-Zn(II) ensembles of experimentally characterized systems (bottom). PDB ID: 1ZE9 (A), 5LFY (B), 2LI9 (C) and 2MGT (D).

The modeling of A*β*-Zn(II) complexes is further complicated by the disordered nature of A*β*.^8^ IDPs lack a well-defined native structure and are thus best described as an ensemble of conformations. Simulating IDPs hence requires classical force fields carefully parametrized to avoid too strong structural propensities favoring folded states.^27,28^ In the last few years, extensive work has dramatically improved the ability of force fields to reproduce experimental data on peptides, unfolded states of proteins and IDPs.^29,29–35^ The remaining inaccuracies in the simulated ensembles can be alleviated using integrative approaches, where experimental data is used as input in the modeling.^36^ One possibility is to restrain simulations to match the experimental data available. In the case of heterogeneous systems, this approach may bias the sampling towards conformations that are not representative of any of the relevant states.^37^ Alternatively, maximum-entropy approaches help generate conformational ensembles that are also compatible with the prior knowledge of the system given, in the case of biomolecular simulations, by the force field.^37,38^ Specifically, Bayesian/Maximum entropy reweighting approaches optimize statistical weights of the snapshots sampled in molecular simulations in order to maximize the agreement with experimental data. These approaches have recently been used in various systems, such as RNA tetranucleotides^39^ or IDP systems.^40,41^

In this work, we focus in the N-terminal region of A*β*, which contains its zinc-binding region.^42^ Using atomistic MD simulations and an optimized force field, we generate conformational ensembles of A*β*-Zn(II) complexes with different coordinations and in different oligomeric states that we have parametrized using the metal center parameter builder package (MCPB.py) developed by Li and Merz.^43^ This approach has recently been used to study complexes of A*β* with Zn(II),^44^ Cu(II) and Al(III).^26^ Here we combine the resulting parameters with the optimized Amber99SB^*⋆*^-ILDN force field, ^45^ which includes corrections for backbone^30^ and sidechain torsions^31^ to the ff99SB force field^46^ that better capture experimental structural propensities in short peptides. In a recent study comparing multiple force fields to study A*β*_42_, this modified force field produced results that compared extremely well against experiments.^47^ We have used these parameter sets to run molecular simulations and validated them against experimental data including NMR chemical shifts and NOEs. Although we find a good agreement between simulation and experiment, we further improve the ensembles using a recently developed Bayesian/Maximum entropy reweighting method.^37^ Our results indicate that A*β*-Zn(II) conformational ensembles may be more heterogeneous than suggested by the NMR models deposited in the Protein Data Bank.

## Materials and methods

### A*β*-Zn(II) models

In order to find minimal model systems for the study of protein-metal interactions, we queried the Protein Data Bank for entries from the full length A*β* protein in complex with Zn(II). The results of this search include structural models of peptides of different variants of human and rat A*β* either as monomers or dimers (see Table 1). Sequence lengths vary between 10 and 16 residues from the N-terminus of A*β* and include all the amino acid residues that have been proposed to interact with the Zn(II) cation.^10,48^ The structures of these peptides have been resolved using solution NMR in the presence of Zn^2+^ in the form of either monomers or dimers, with one or two cations per complex. All models have different ligand combinations (see Figure 1). In all four cases, NOE restraints are available at the NMR Restraints Grid, and for all dimers chemical shifts are also available at the Biological Magnetic Resonance Database (BMRB).

**Table 1:**
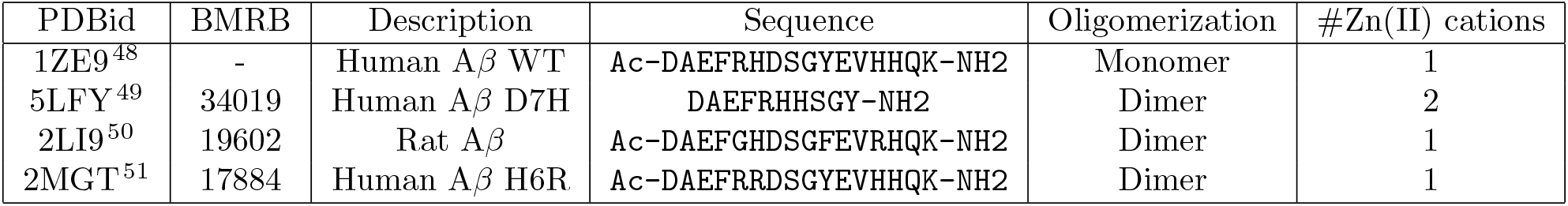
Molecular systems included in this study

### Parametrization of protein-metal interactions

All bonded models used have been parametrized using the MCPB.py tool by Merz and co-workers,^43^ which is distributed with Ambertools^1^. We performed all DFT quantum mechanical calculations using the B3LYP/6-31G*^52,53^ method in the Gaussian16 package.^54^ For each of the model systems, generating new parameters requires three different steps: geometry optimization, estimation of force constants using the Seminario method^55^ and calculation of point charges using the restrained electrostatic potential (RESP) model. ^56^

Three out of four of the systems we are studying are protein dimers, where two exact copies of the same protein chain are coordinated to either one or two Zn(II) cations. Due to minor discrepancies between dimers present in the experimental structures, we have obtained slightly different values of equilibrium geometry parameters, force constants, and charges for the atoms in each of the chains. To make the two chains in the dimers indistinguishable in our simulations, the protein-metal force field parameters obtained should be exactly the same. We have followed the approach taken by Peters et al. when parametrizing the ZAFF force field^21^ and averaged the charges, equilibrium geometry parameters and force constants of bonds, angles and dihedrals of all duplicate values. Thereby, we have obtained the same parameter sets for both protein chains present in dimer systems.

All the files used to parametrize the bonded models and the resulting parameters are available at https://osf.io/y4zk5/.

### Molecular dynamics simulations

We have inserted all systems within cubic boxes and solvated them with TIP3P water molecules.^57^ We added a 0.1 M concentration of Cl^−^ and Na^+^ ions to all systems, and also neutralized the total charge of the system when necessary. The systems were energy-minimized using the steepest descent algorithm and then equilibrated in two stages. First, we run a 100 ps simulation in the NVT ensemble and then another 100 ps in the NPT ensemble, both including position restraints in the protein heavy atoms. Molecular Dynamics (MD) simulations were performed in the NVT ensemble, using an integration time step of 2 fs. Constant temperature and pressure were set by coupling the system to a Parrinello-Rahman barostat at 1.0 bar and a velocity-rescaling thermostat at 278 K,^58–60^ corresponding to the temperature of the NMR experiments. Equilibrium simulations were run for 5-10 *μ*s, depending of the size of the system and convergence of the simulations, with the Amber99SB^*⋆*^-ILDN force field^30,31^ using the Gromacs software package (version 2020).^61^

### Analysis, validation, and reweighting

To analyze the results of the simulations we have used a combination of Gromacs tools and in-house scripts using the MDtraj Python library. ^62^ We have validated the results of our simulations against experiments back-calculating a series of NMR observables from the simulations. We used the SPARTA+ program to back-calculate chemical shifts.^63^ Additionally, we estimate distances corresponding to the observed NOEs using *r*^−6^ averaging.^64–66^

We additionally perform a Bayesian/Maximum entropy reweighting using the BME program^37^ to refine simulations using the experimental data available, in this case NOE distance restraints. The main goal of the reweighting of simulations is to obtain an optimized set of statistical weights for the simulation ensemble that are in better agreement with experiments given a forward model for the experimental observable.^36^ More information about BME is provided in the Supporting Information.

## Results and discussion

### Molecular simulations with new force fields parameters

In a first attempt to model the complexes of A*β*-Zn systems, we used non-bonded models using the recently optimized Amber99SB-disp force field.^34^ Unfortunately, these efforts resulted in the loss of the tetrahedral coordination of the metal within a few hundreds of nanoseconds (see Supporting Information S1-S5 for additional details). This result adds to the mounting evidence that non-bonded models cannot be used to appropriately model A*β*-metal complexes.^11^ We hence resorted to generate parameters for the metal using a bonded model. Specifically, we used the MCBP.py package to generate parameters for all the A*β*-Zn(II) complexes in Table 1 (see Methods).

Using the new parameters, we have run 5 *μ*s atomistic MD simulations for all four systems. This amount of simulation time seems sufficient to converge the distribution of *R*_*g*_ for the monomer (1ZE9) and D7H mutant dimer (5LFY), which remains stable for over half the simulation runs (see Fig. S6 in the Supporting Information). In the case of the rat and H6R mutant, we extended the simulation runs up to 10 *μ*s in order to sample conformational space more exhaustively. In Figure 2 we show the values of the C^*α*^ − *RMSD* and the radius of gyration (*R*_*g*_) for each of them. The *RMSD* values typically remain under 5 Å for most of the simulation time. However, with the exception of the D7H mutant dimer (5LFY), we observe relatively large fluctuations that suggest that the simulations explore a variety of conformations that are dissimilar from those reported in the PDB. On the other hand, we find that the *R*_*g*_ values remain stable throughout the simulations, although two of the systems are notably more compact than the PDB models. This effect is likely due to the influence of the TIP3P water model used in our simulations, which is known to produce these effects.^67^

**Figure 2:**
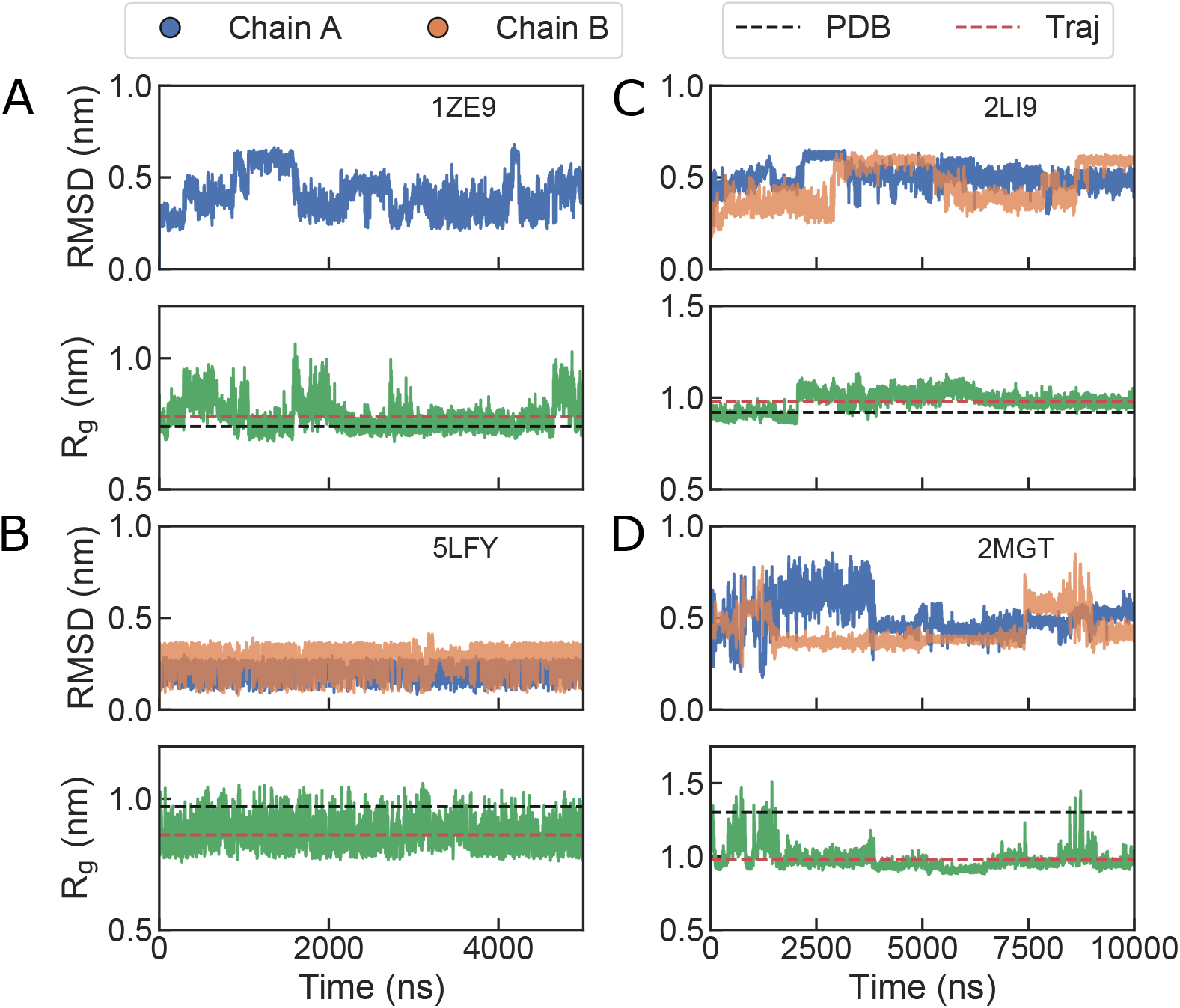
*RMSD* (A) and *R*_*g*_ (B) values calculated from the 5 *μ*s simulations using the parametrized bonded models. Mean *R*_*g*_ values calculated from PDB models are shown by black lines and mean *R*_*g*_ values calculated from our simulations are shown by red lines.

In order to identify the regions of the protein with the largest fluctuations, we also report values for the *RMSF* for all systems (see Figure 3A). As expected, the highest *RMSF* values correspond to regions at the N and C-termini of each system, except in the case of the D7H mutant dimer (5LFY). In this case, the low *RMSF* values in the N-termini are related to the coordination of the Zn^2+^ cation by the oxygen from the carbonyl of the first amino acid in the peptide sequence. On the contrary, in the case of the monomer (1ZE9), H6R mutant dimer (2MGT) and rat A*β* dimer (2LI9), fluctuations are on average larger and their values are greatest at the N and C-termini.

**Figure 3:**
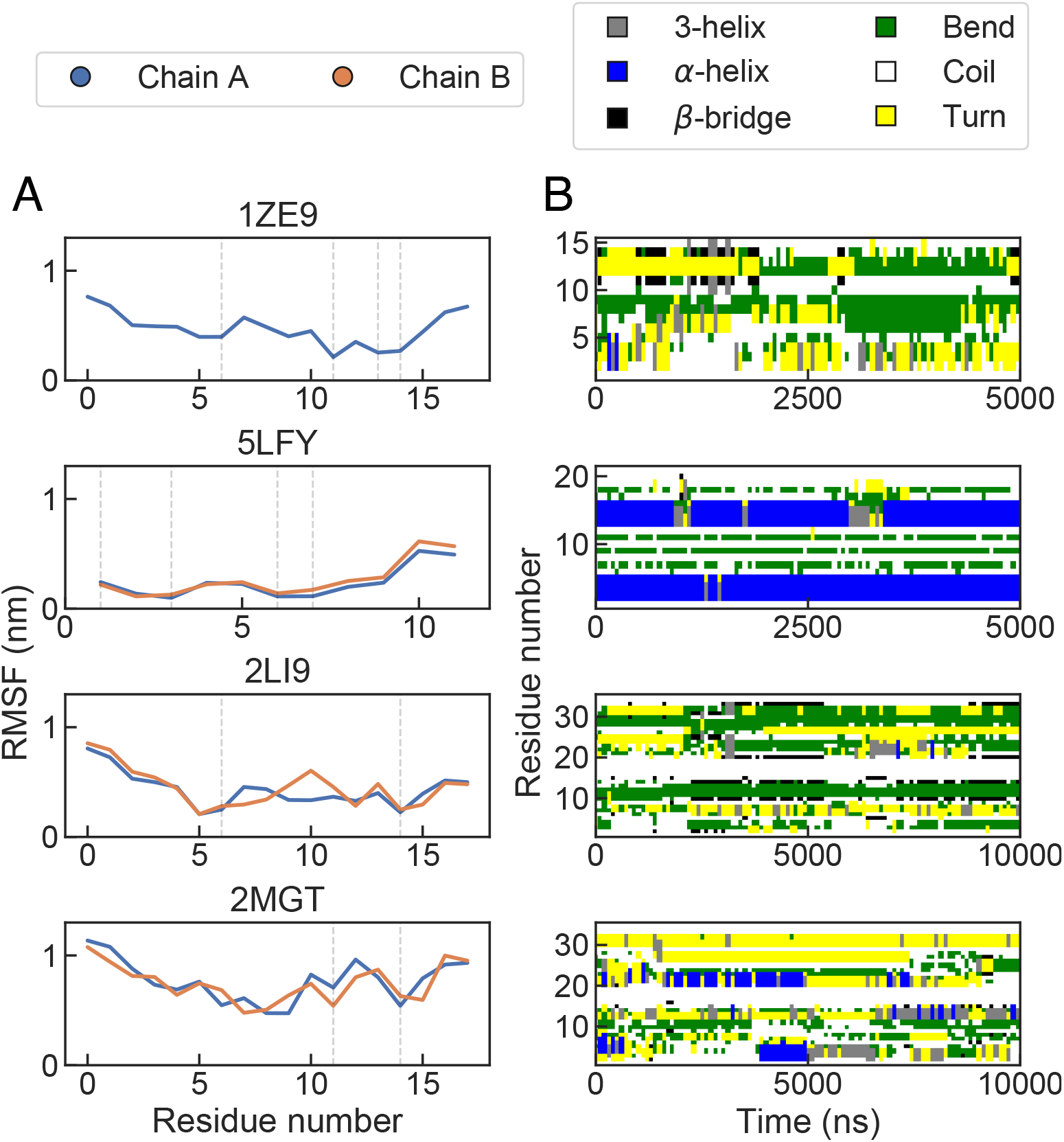
(A) Per-residue *RMSF* values calculated for each of the A*β*-Zn systems. For dimers, we represent each chain with a different colour. Vertical dashed lines mark the Zn-chelating residues. (B) Secondary structure assignment calculated with DSSP.

The trends observed in the *RMSF* are consistent with the very different structural propensities in the metal-bound peptides. In Figure 3B we show the assignments to different types of secondary structure obtained using the DSSP algorithm. ^68^ The regions that more stably keep the secondary structure coincide with those with the lowest *RMSF* values, which also contain the chelated residues. This behavior is clearest for the D7H mutant dimer (5LFY), where the *α*-helices remain fully formed in both peptide chains through almost the complete duration of the simulation trajectory.

In order to visualize the differences between the ensemble of conformations sampled during the simulations and the NMR structures, we have calculated the contact maps, which we compare with the averages from the 20 structural models deposited in the PDB for each of our systems (see Figure 4). We find that despite the intermediate values of the *RMSD* reported above, the contact maps from the simulations are largely consistent with those of the experimental models. Specifically, close contacts from the NMR ensembles are mostly recapitulated by the simulation results. This is expected because the conformational dynamics in our complexes are restricted by the interactions with the metal cations.

**Figure 4:**
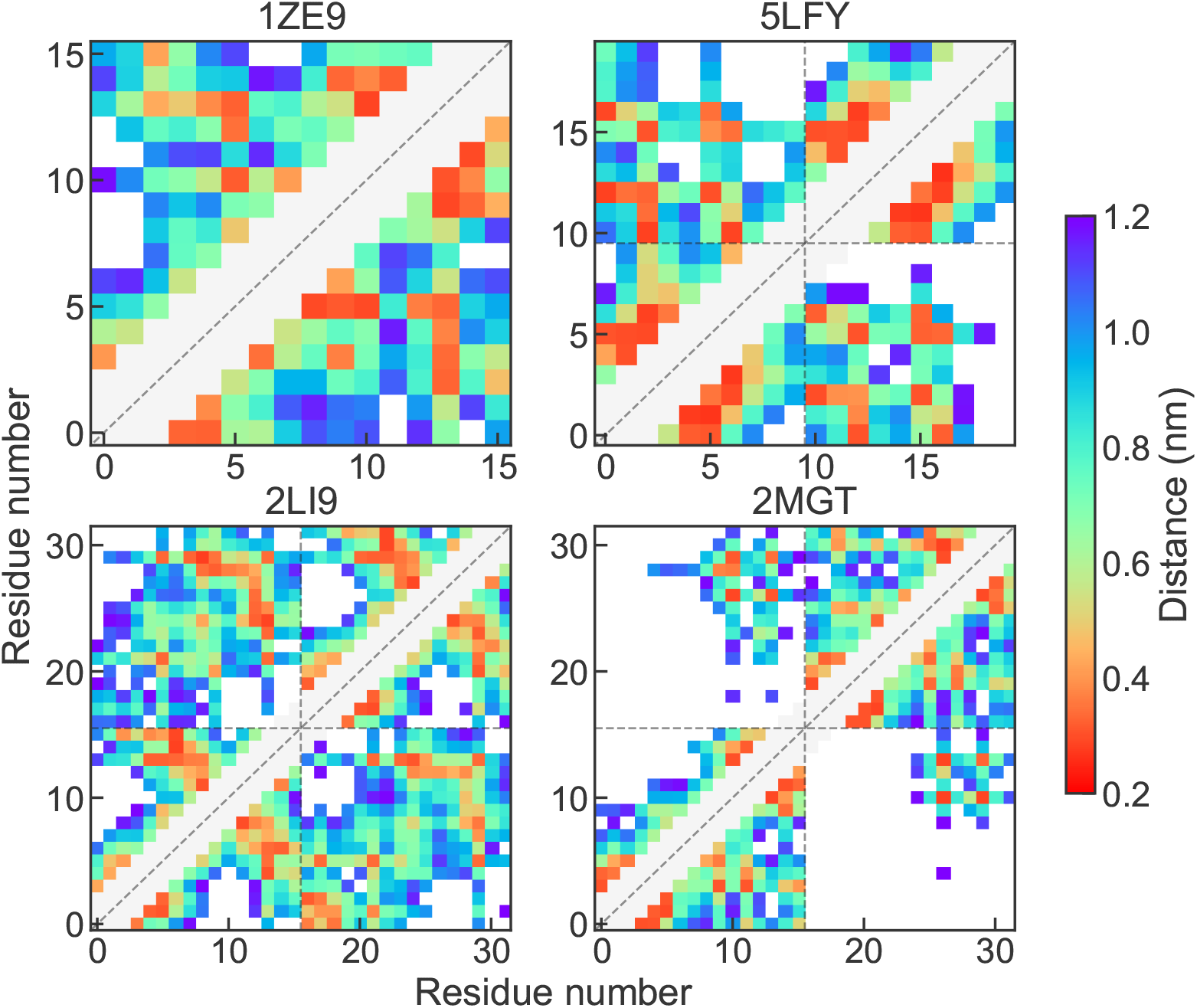
Contact maps calculated from PDB structures (lower triangle) and MD trajectories (upper triangle).

### Validation against NMR data

In order to assess the validity of our simulations more carefully, we compare the results obtained with the combination of the new metal parameters with the Amber99SB^*⋆*^-ILDN force field directly against available experimental NMR data. First, we compare experimental chemical shifts with values back-calculated from the simulations using SPARTA+^63^ (see Methods). In Table 2 we report the *RMSD* and *χ*^2^ values of the calculated chemical shifts, and compare them with those for the NMR models. Overall, both the *RMSD* and *χ*^2^ values obtained from the 20 structures deposited in the PDB and our 5-10 *μ*s simulations are in the same range, indicating that our simulations have reached a comparable level of accuracy as the experimental models for the A*β*-Zn(II) systems.

**Table 2:**
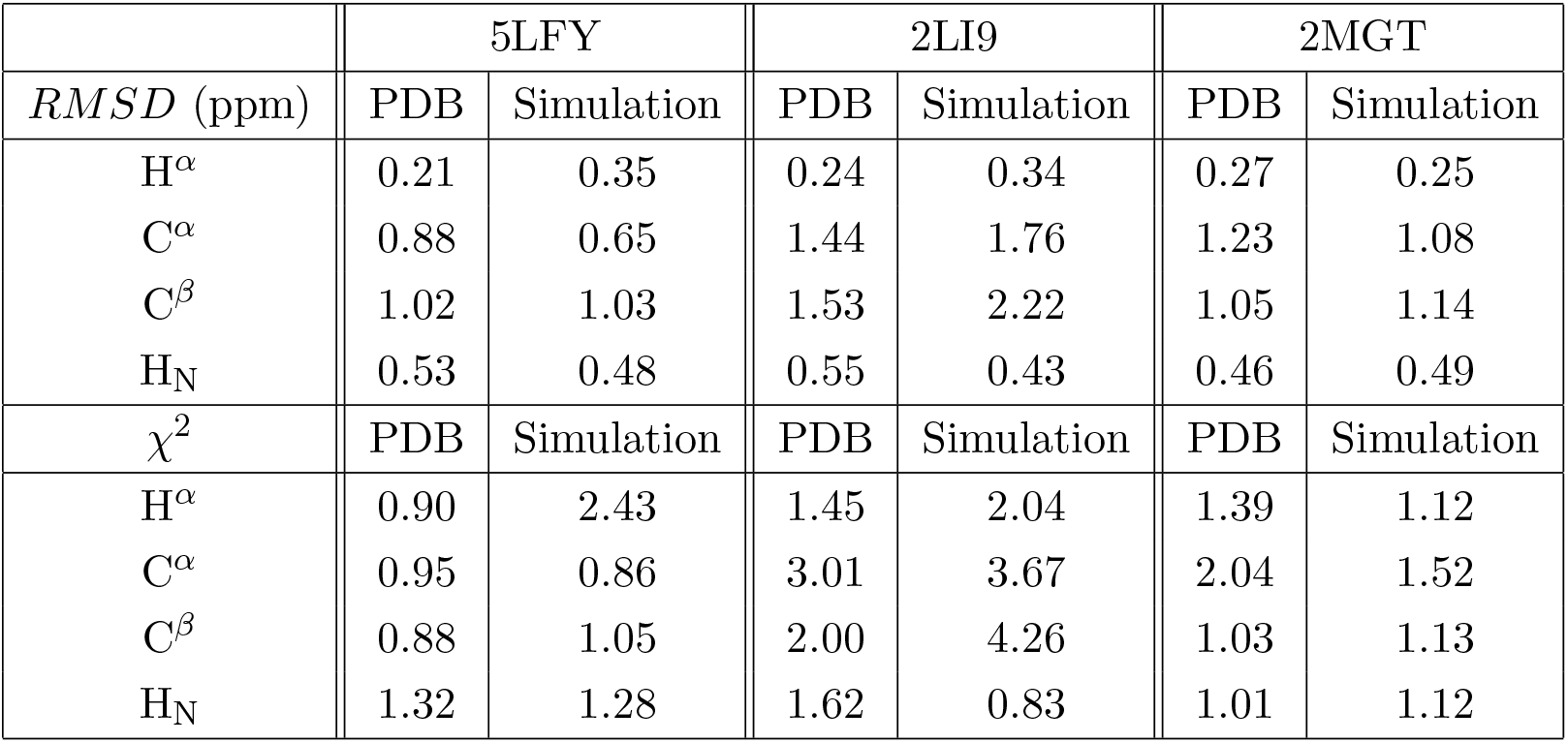
Calculated *RMSD* and *χ*^2^ values of chemical shifts back-calculated from PDB structures and simulations.

There are, however, some noteworthy inconsistencies between the calculated chemical shifts and the experimental measurements (see Figure 5). For example, the simulations do not completely track the overall tendency of the chemical shift of H*α* for residues 1 and 3 of the D7H mutant (5LFY model). In the case of C^*α*^ and C^*β*^ chemical shifts, the agreement is worst for the rat peptide (2LI9), although the errors are also largest for the models deposited in the PDB and the overall trends are captured correctly. Instead, for both the D7H and H6R mutants (5LFY and 2MGT, respectively) the agreement with experiments is excellent for the carbon chemical shifts. Lastly, for the HN protons, the simulations for the rat peptide (2LI9) result in the best agreement obtained shown both from *RMSD* and *χ*^2^ values.

**Figure 5:**
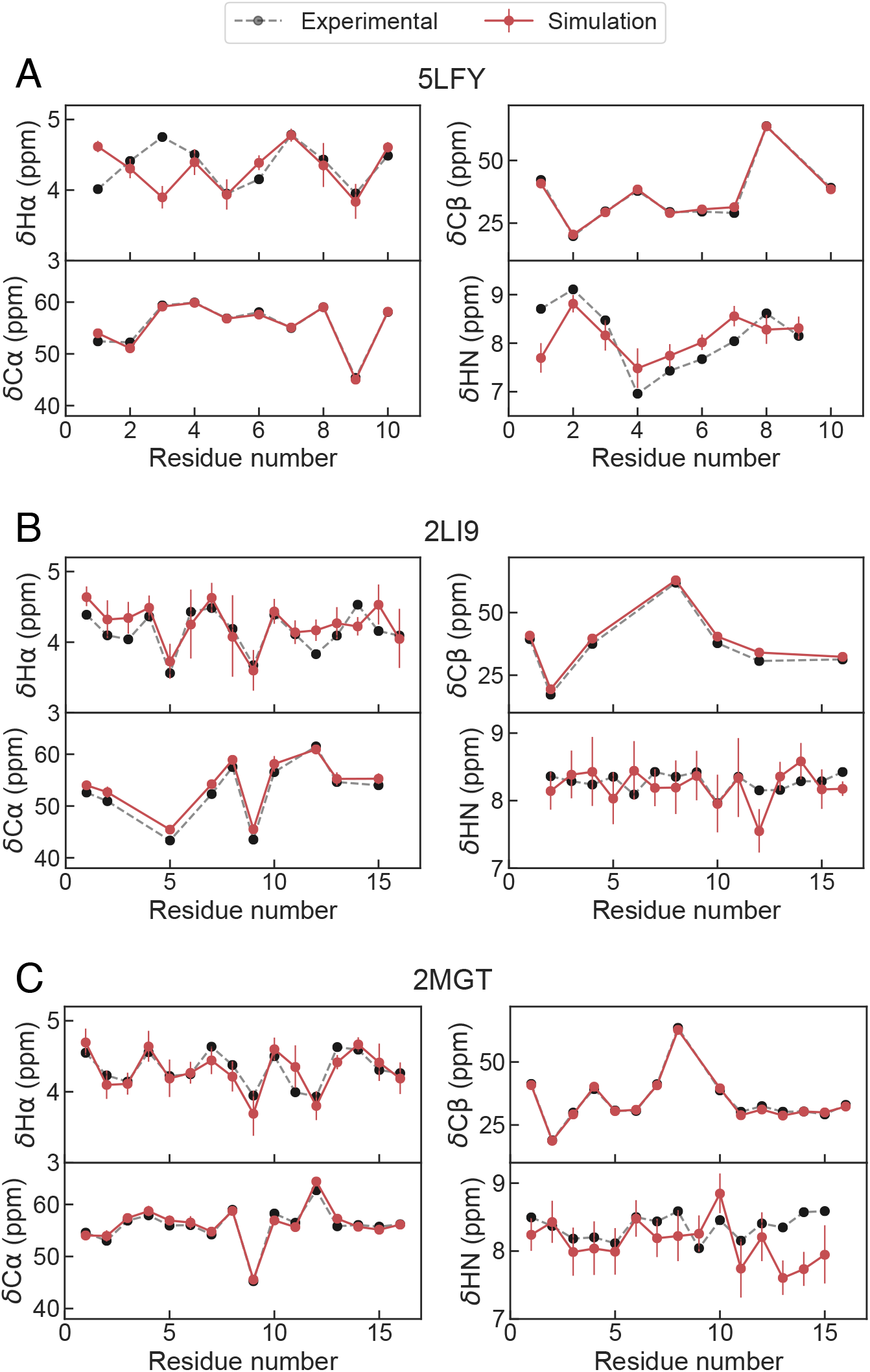
Chemical shifts calculated for the 5LFY (A), 2LI9 (B) and 2MGT (C) systems with experimental data available. Experimental values are shown in black and back calculated chemical shifts are shown in red.

Additionally, we have calculated average distances from the simulations for all the atom pairs involved in measurable NOEs (see Figure 6). We note that for all the dimers, the intra-chain NOE distance restraint values are duplicated for chains A and B in the BMRB entries (see Figure S7). To estimate the average distances from the simulations, we assume that the chains are indistinguishable, as they are in the experiment, and keep only one set of NOE values. Accordingly, we report a single set of distances averaging distances from both chains. When no errors were reported, we arbitrarily assign a 20% error to the measured value.

**Figure 6:**
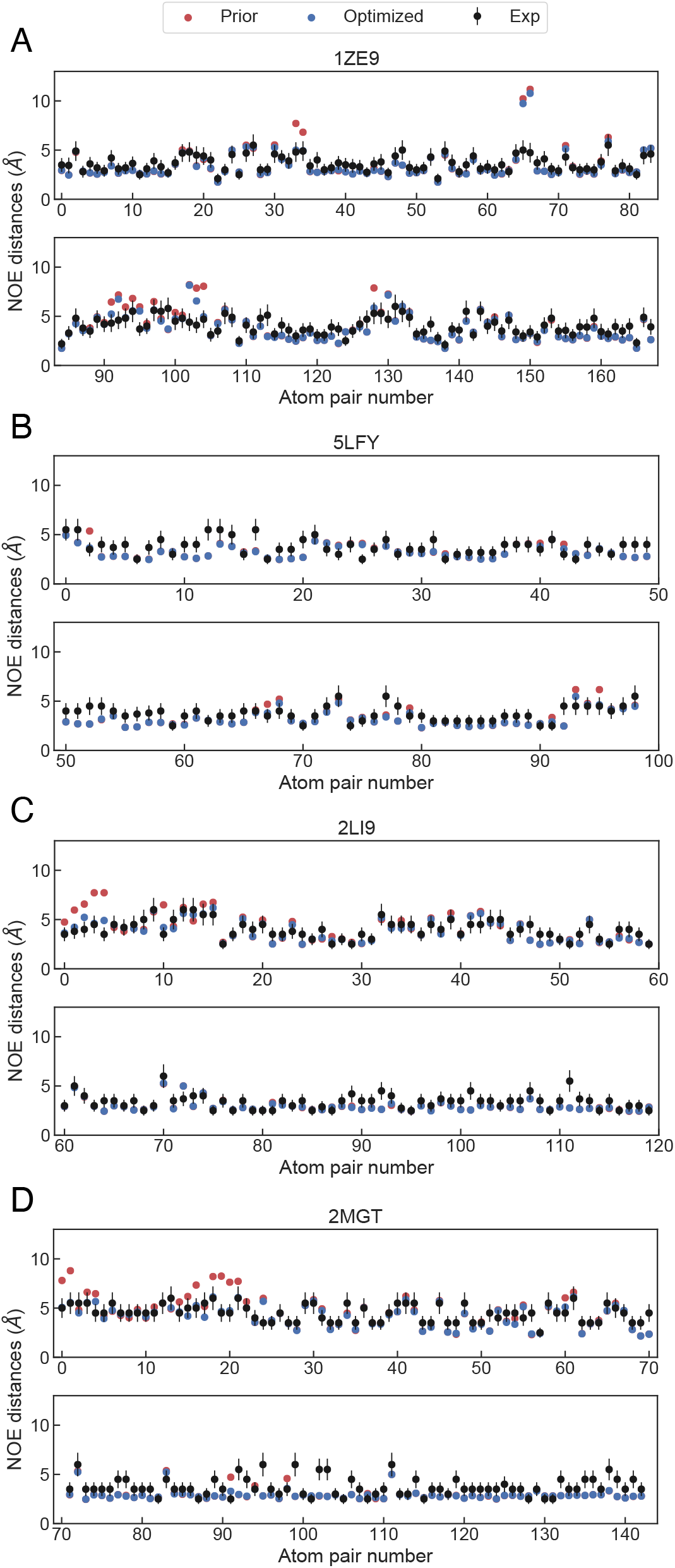
NOEs calculated for the the 1ZE9(A), 5LFY (B), 2LI9 (C) and 2MGT (D) systems with experimental data available. Experimental values are shown in black, back-calculated NOEs are shown in red and NOEs calculated after optimization are shown in blue.

Overall, the simulations with the new bonded parameters and the Amber99SB^*⋆*^-ILDN force field successfully capture the vast majority of the experimental NOE distances. Interestingly the *χ*^2^ value obtained for each system is 0.078 for the monomer (1ZE9), 0.076 for the D7H mutant (5LFY), 0.081 for the rat dimer (2LI9) and 0.083 for the H6R mutant (2MGT). Nonetheless, some values of NOE distances are overestimated from the simulations e.g. distances 65 and 66 of the monomer (1ZE9), distances 0 to 10 in the rat dimer (2LI9) or distances from 15 to 20 of the H6R mutant (2MGT) (see Figure 6).

### BME reweighting improves agreement with NMR data

The overestimation of some NOE distances from the unbiased simulations suggests there is room for improvement in the conformational ensembles. Therefore, we have reweighted our simulations to further improve the quality of the ensembles obtained for the systems using the BME program developed by Bottaro et al.^37^ which relies on Bayesian/Maximum entropy approach (see Methods) using the NOE data from Figure 6. Ensemble reweighting helps improve the agreement with experiment for all systems, significantly in some cases, as indicated by a decrease in *χ*^2^. After optimization the *χ*^2^ values are of 0.050 for the monomer (1ZE9), 0.045 for the D7H mutant (5LFY), 0.046 for the rat dimer (2LI9) and 0.032 for the H6R (2MGT), a 36%, 55%, 43% and 61% improvement, respectively.

In order to illustrate the effect of the reweighting, we show the distribution of *R*_g_ calculated for all systems (see Figure 7). In the case of the monomer system (1ZE9) a bimodal distribution of the *R*_g_ is obtained. Before reweighting, the distribution is showing that contorted conformations are much more populated, while the reweighting reduces that bias. In Figure 7B, we show selected snapshots corresponding to the low and high *R*_*g*_ peaks. Similarly, in the case of the rat dimer (2LI9), the force field prior distribution indicates a greater propensity for extended conformations, which is modulated by the reweighting. As for the monomer, we show snapshots corresponding to both populations in Figure 7C. The effect the reweighting in the case of the H6R mutant (2MGT) is more modest, but again shows that ensembles including extended conformations are in better agreement with experiment. These results clearly show the effect the BME reweighting has over simulations and proves that, generally, the most accurate ensemble description of A*β*-Zn(II) systems are built with more hetereogeneous ensembles than would be expected from the NMR models alone.

**Figure 7:**
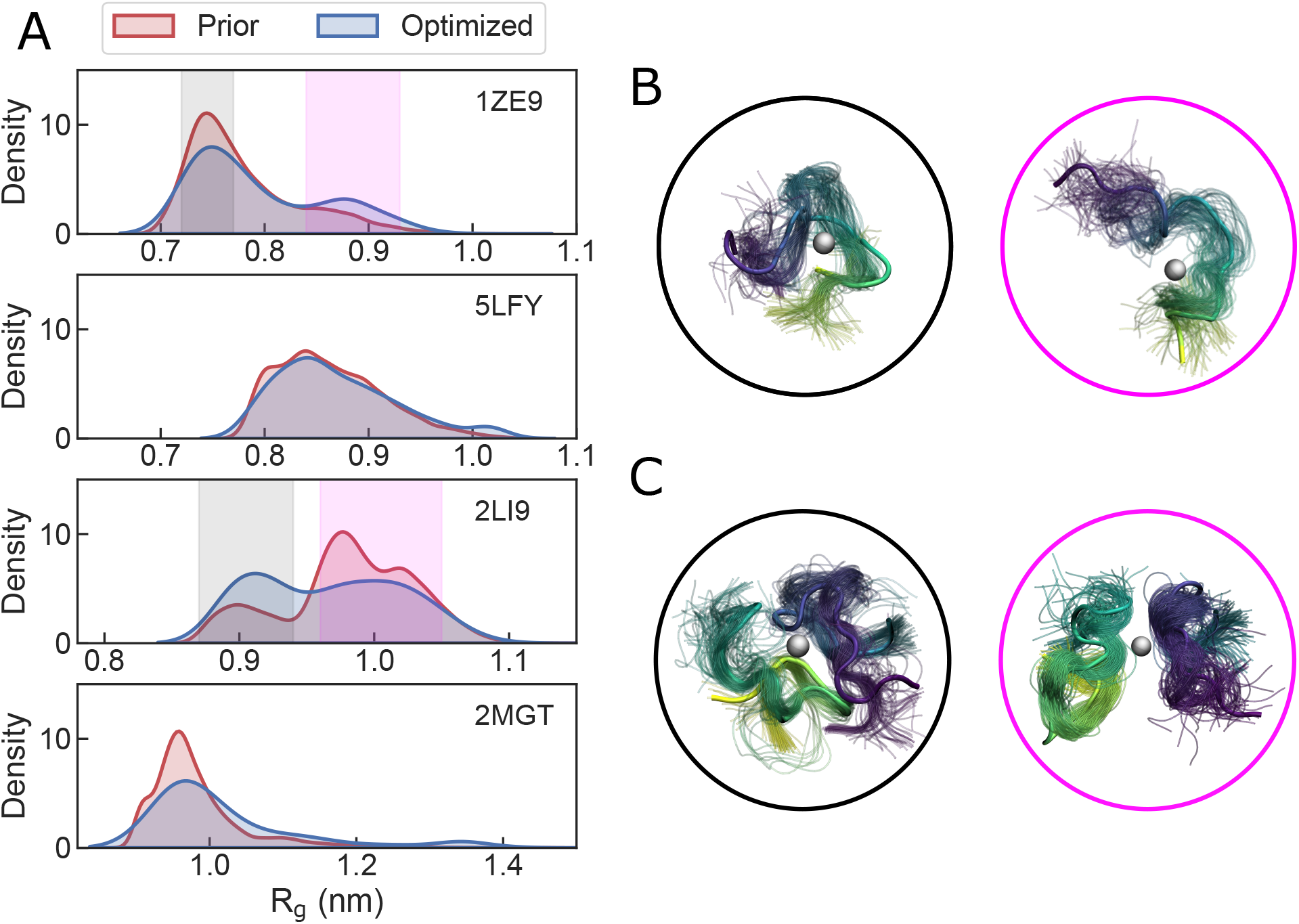
Distribution of R_g_ calculated from the prior distribution and optimized distribution (A), different R_g_ conformers populated by the 1ZE9 model (B) and 2LI9 model (C), where contorted conformers are shown in black circles and extended conformers in magenta circles.

## Conclusions

In this work, we have produced bonded Zn(II) models of four A*β*-Zn(II) systems for the Amber99SB^*⋆*^-ILDN force field. We note that other force field-water model combinations may further improve the accuracy of the results. Using these parameters, we have run 5-10 *μ*s simulations of four different systems and validated the results against experiments. The simulations have sampled well beyond the conformational space of the models reported in the PDB as shown by the *RMSD* of C^*α*^ and contact maps. This extended sampling nevertheless compares well with experimentally measured chemical shifts and NOEs.

In order to improve the agreement between simulations and experiments, we have reweighted our simulations using the BME program^37^ using NOE data. Interestingly, the improvement of the simulations shows that the best agreement is obtained when both collapsed and extended conformations of the systems are present in the conformational ensembles. This may have implications for our understanding of the role of Zn(II) in the aggregation of A*β*, which has been explored in the past using the more structured NMR models as initial structures for MD simulations.^10^

We hope that the parameters we have produced for all four systems will help on gaining a better understanding of A*β*-Zn(II) interactions. The parameters that we make publicly available for all four complexes will afford more accurate simulations and may be applied in studies of primary and secondary nucleation of fibrils.^69^

## Acknowledgement

Financial support comes from Eusko Jaurlaritza (Basque Government) through the project IT1584-22 and from the Spanish Ministry of Science and Universities through the Office of Science Research (MINECO/FEDER) through grant PGC2018-099321-B-I00. DDS acknowledges the Spanish Ministry of Science and Universities for a Ramón y Cajal contract (Grant RYC-2016-19590). The authors thank Sandro Bottaro for help using the BME program and Kresten Lindorff Larsen for useful discussions. The authors also acknowledge the staff at the DIPC Supercomputing Center for technical support.

## Data and Software Availability

All the input files required to reproduce our results, together with structure and parameter files required to run MD simulations on our systems are supplied as Supporting Information.

## Supporting Information Available

Supplementary Methods and eight additional figures.

## Footnotes

1 https://ambermd.org/AmberTools.php

