## Supporting Information for "Atomistic molecular simulations of A*β*-Zn conformational ensembles"

### Supporting Information: Atomistic molecular simulations of A$\beta$-Zn conformational ensembles

#### Simulations with nonbonded models of Zn(II)

To test the performance of nonbonded models we tested the Amber99SB-disp with its default Zn(II) model.[S1] Amber99SB-disp is one of the most recently parameterized force fields to simulate disordered proteins. We ran the simulations with the same specifications as described in the manuscript (see Methods).

Interestingly, as soon as the minimization process was performed, Zn(II) was found coordinating to two additional water molecules in an octaherdal arrangement. Although Zirah et al. report that in the 1ZE9 model, Zn(II) coordinates A$\beta$ through His6, Glu11, His13 and His 14,[S2] during the equilibration all histidines dissociate from Zn(II) and additional water molecules substitute them in an octahedral arrangement (see Figure S1).

Although coordination of histidines is kept for the 2LI9 model (see Figure S3), additional water molecules are found to coordinate Zn(II) as well as Ser8 and Gly9 through its carbonyl group, once again in an octahedral arrangement (see Figure S4). Coordination with glutamic/aspartic acids are kept throughout all simulation timeseries, which suggests that nonbonded models favour electrostatic interactions over orbitalic interactions.

In conclusion, evidence points out that nonbonded models fail at accurately represeting His-Glu/Asp mixed metal centre coordination as well as tetrahedral arrangements of said metal centres.

#### Convergence of simulations

To check the convergence of our simulations we have calculated cumulative averages of radius of gyration ($R_g$) values in ten intervals. After 5$\mu$s, simulations of the 1ZE9 and 5LFY models are well converged (see Figure S6A and S6B). On the other hand, for the 2LI9 and 2MGT, 5 additional $\mu$s were needed, simulations being extended up to 10$\mu$s (see Figure S6C and S6D).

#### Reweighting of the simulations using NOEs

The Bayesian/Maximum Entropy method (BME) allows reweighting the frames in a molecular dynamics (MD) trajectory to improve agreement with experiment.[S3,S4] In practice, the goal is to obtain an optimized set of statistical weights for the simulation frames, $w_j$, by minimizing the negative log posterior function

$$\mathcal{L}(w_1...w_n) = \frac{1}{2}\chi^2(w_1...w_n) - \theta S_{rel}(w_1...w_n) \tag{1}$$

where, $\chi^2$ quantifies the agreement between the experimental ensemble averages of an observable and the corresponding values calculated from the reweighted simulation ensemble.

In the equation above, the relative entropy $S_{rel}$ is expressed as

$$S_{rel} = \sum_{j}^{n} w_j \log\left(\frac{w_j}{w_j^0}\right) \tag{2}$$

where the sum runs over the number of frames, $n$. $S_{rel}$ measures the deviation between the original weights, $w_j^0$ ($1/n$ in the case of having performed unbiased MD) and the optimized weights. Lastly, the global parameter $\theta$ in Equation 1 acts as a "pseudo-temperature" that determines the relative weights of $\chi^2$ and $S_{rel}$ in the optimization. Low values of $\theta$ will enforce a strong agreement with experiment, while large $\theta$ will favour the original unweighted ensemble. In addition, $\theta$ also determines the number of effective frames taken from the forward model that effectively contribute to the calculated averages, $N_{eff} = \exp(S_{rel})$.

In order to perform a BME reweighting, an adequate $\theta$ value was selected. To do so, we looked at how the $\chi^2$ and $N_{eff}$ values changed for different $\theta$ values. We looked for the best compromise between having a significant number of effective frames and an acceptable $\chi^2$ reduction (see Figures S8, S10, S12, S14).

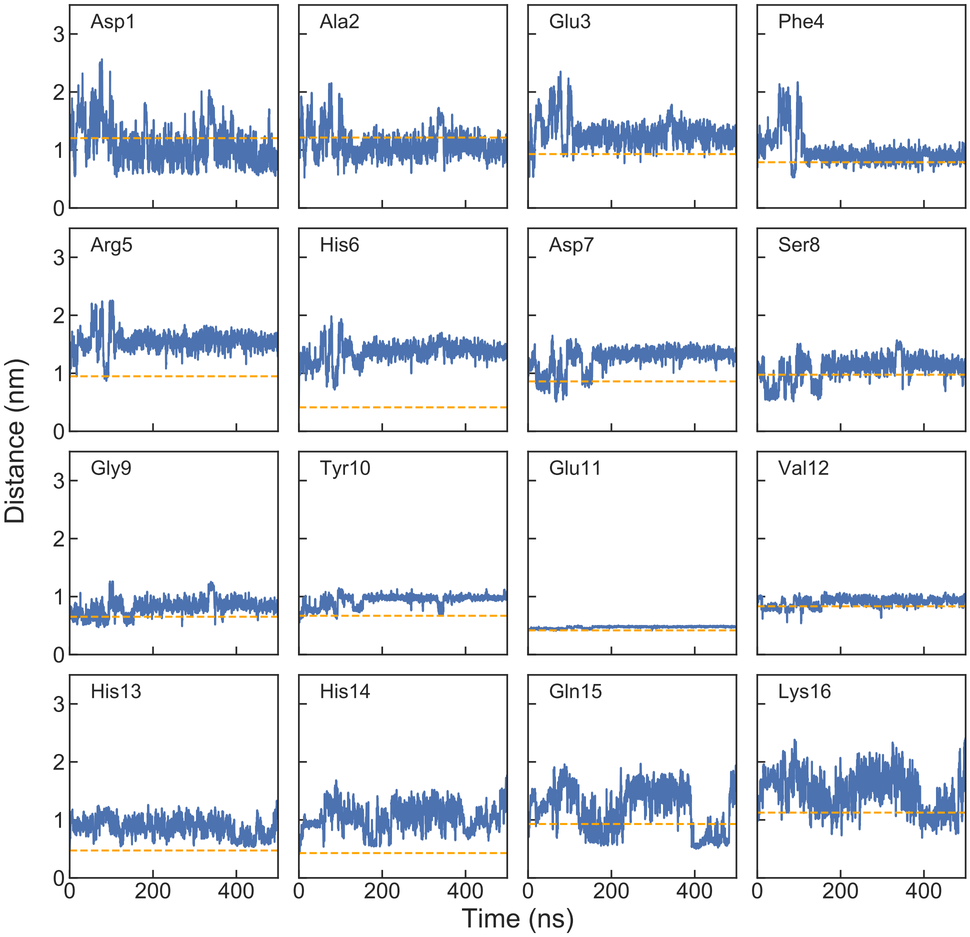


Figure S1: Every residue-Zn(II) distances calculated for the simulation with the 1ZE9 model and Amber99SB-disp force field. Reference distances taken form the PDB are shown with an orange line.

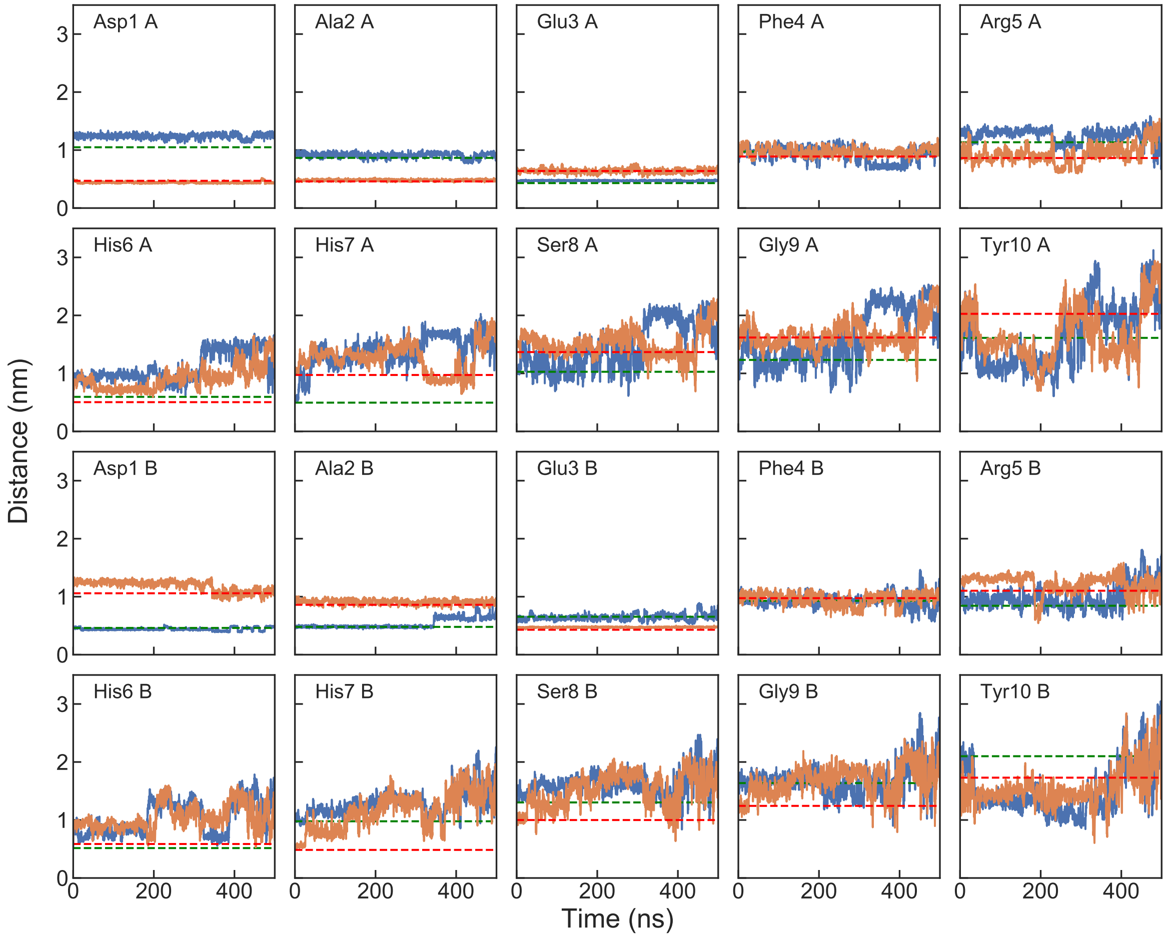


Figure S2: Every residue-Zn(II) distances calculated for the simulation with the 5LFY model and Amber99SB-disp force field. Reference distances taken form the PDB are shown with a green line for Zn1 and in red for Zn2, where Zn1 coordinates Asp1$^{A}$, Glu3$^{B}$, His6$^{A}$ and His7$^{B}$ and Zn2 coordinates Asp1$^{B}$, Glu3$^{A}$, His6$^{B}$ and His7$^{A}$.

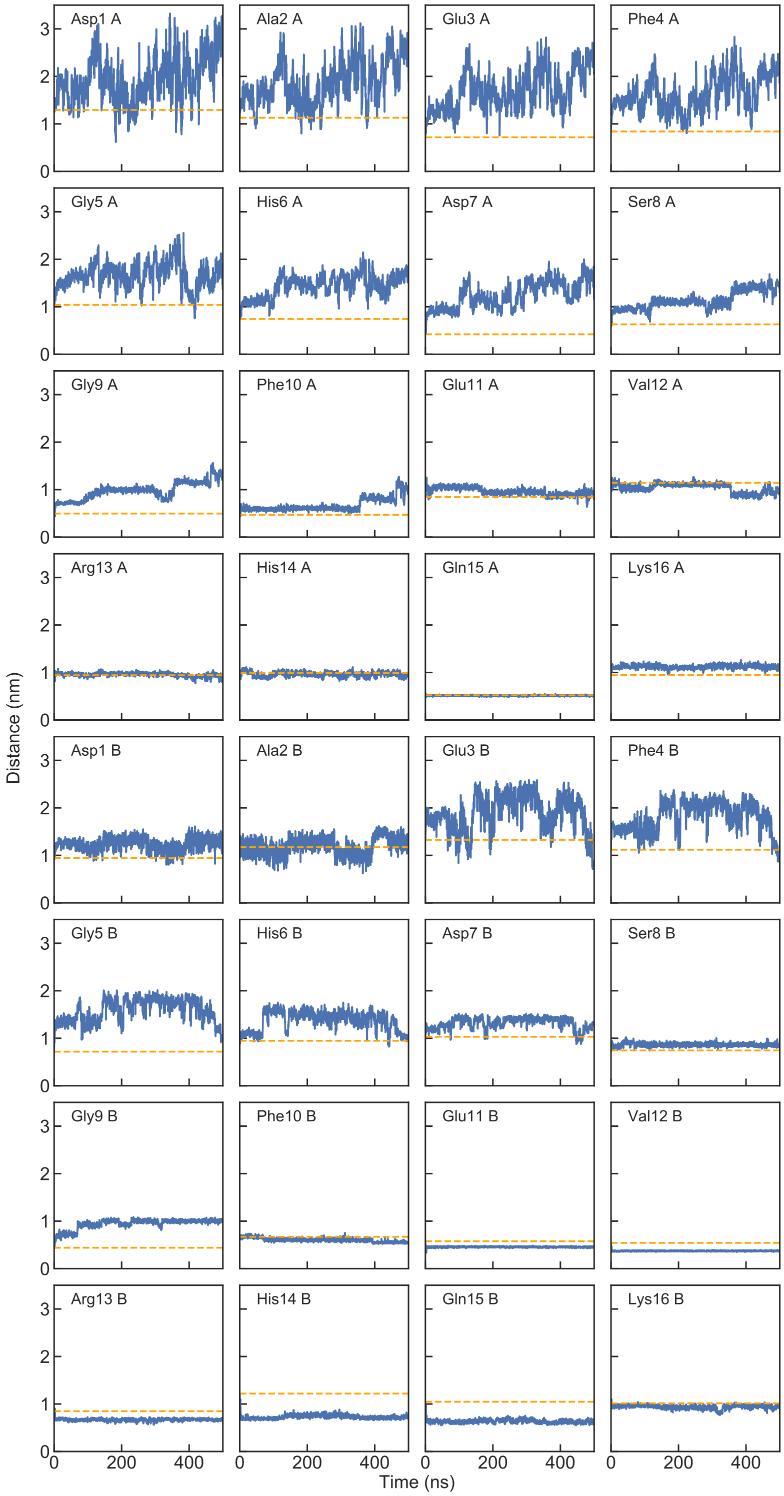


Figure S3: Every residue-Zn(II) distances calculated for the simulation with the 2LI9 model and Amber99SB-disp force field. Reference distances taken form the PDB are shown with an orange line.

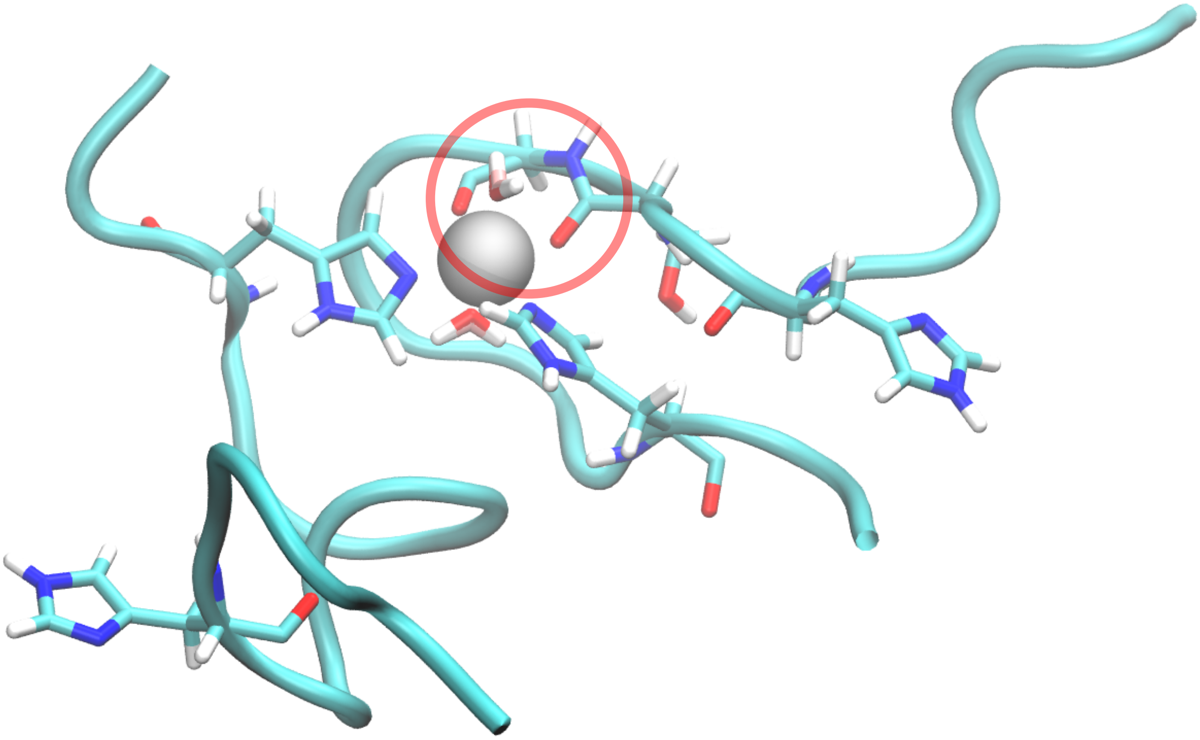

Figure S4: Snapshot that shows coordination of the 2LI9 model with the Amber99SB-disp force field. Carbonyl group of Ser8 and Gly9 are highlighted in the red circle.

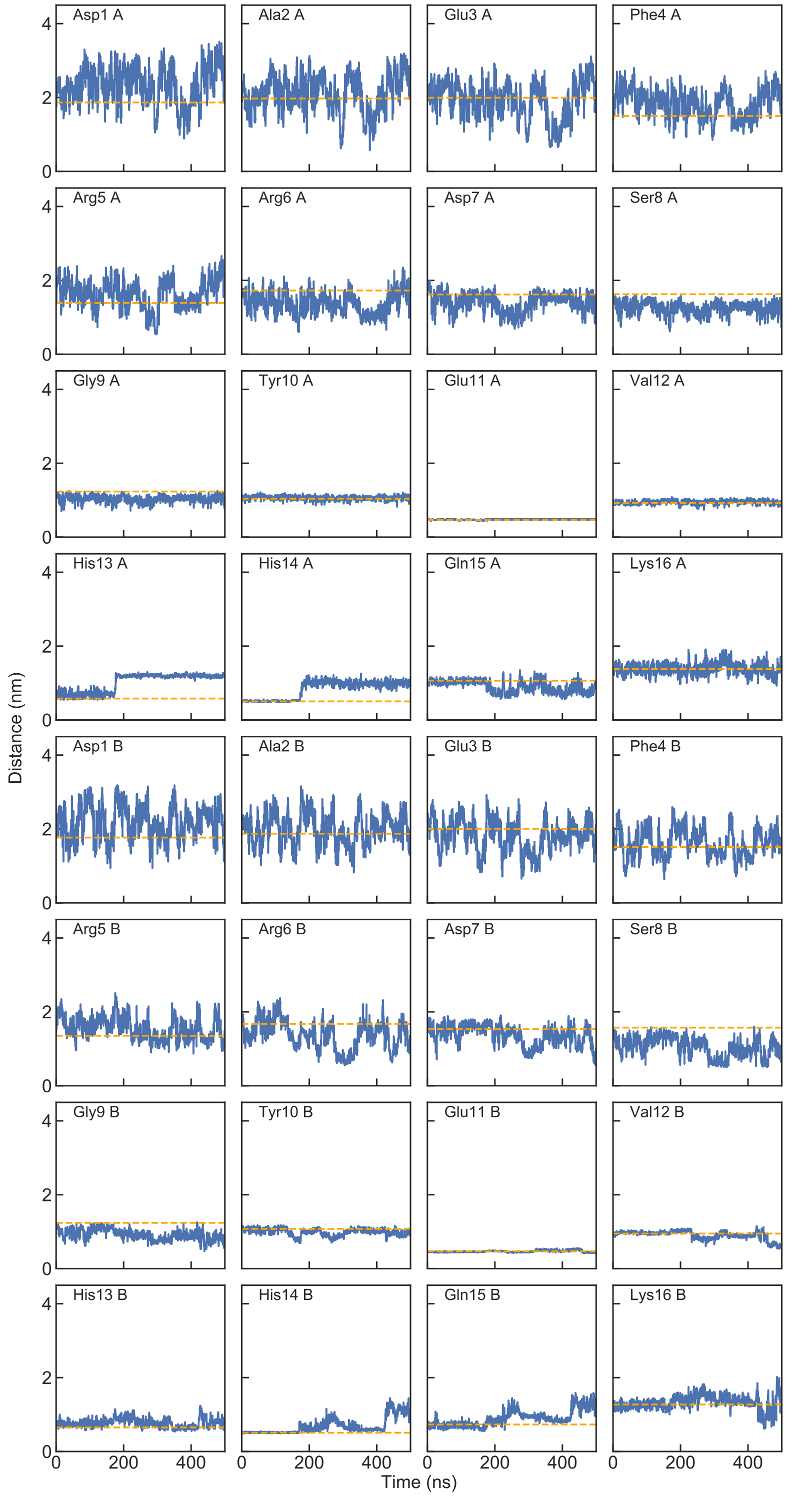


Figure S5: Every residue-Zn(II) distances calculated for the simulation with the 2MGT model and Amber99SB-disp force field. Reference distances taken form the PDB are shown with an orange line.

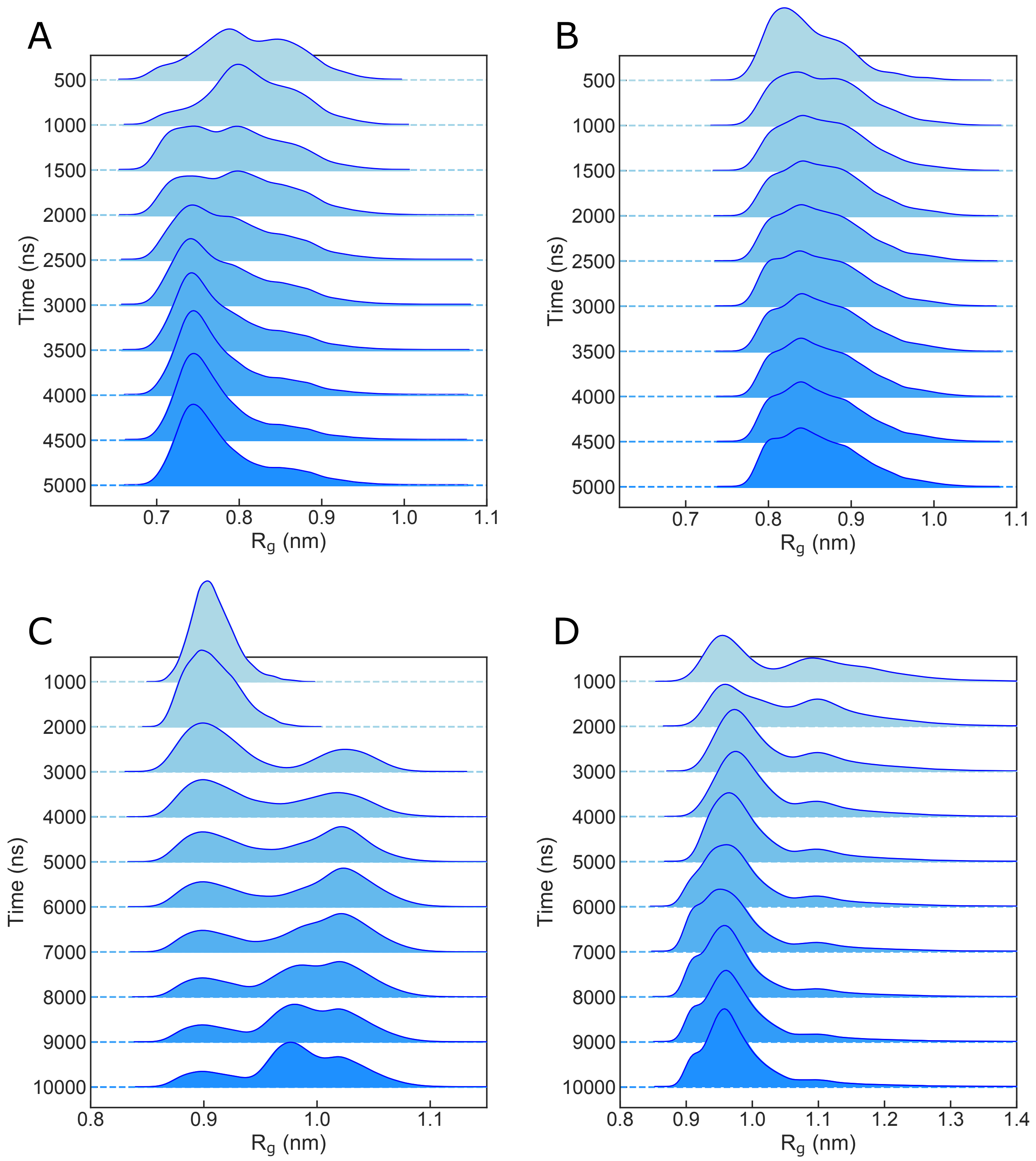


Figure S6: Cumulaive averages of $R_g$ calculated for the 5$\mu$s simulation of 1ZE9 (A), 5$\mu$s simulation of 5LFY (B), 10$\mu$s simulation of 2LI9 (C) and 10$\mu$s simulation of 2MGT (D) with the Amber99SB*-ILDN force field and bonded model of Zn(II).

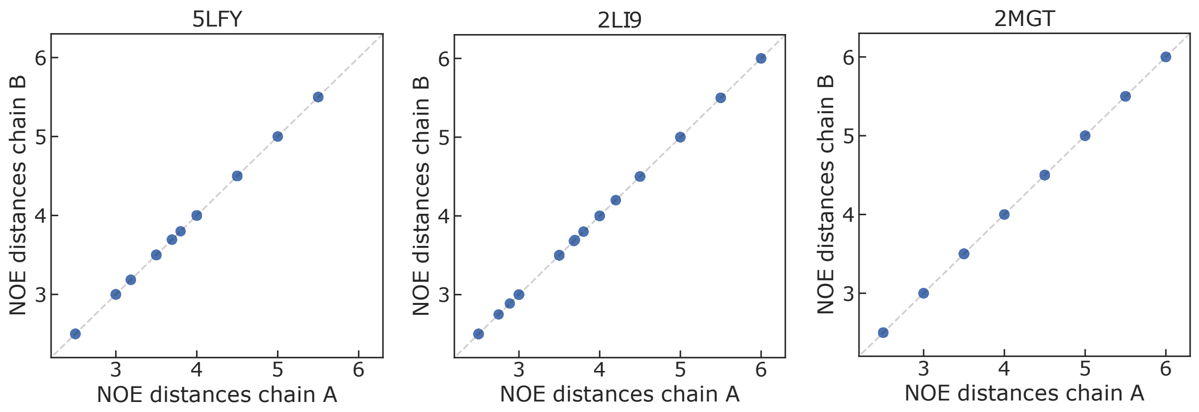


Figure S7: Correlation between experimental NOE distances reported for chain A and chain B of the three dimer systems.

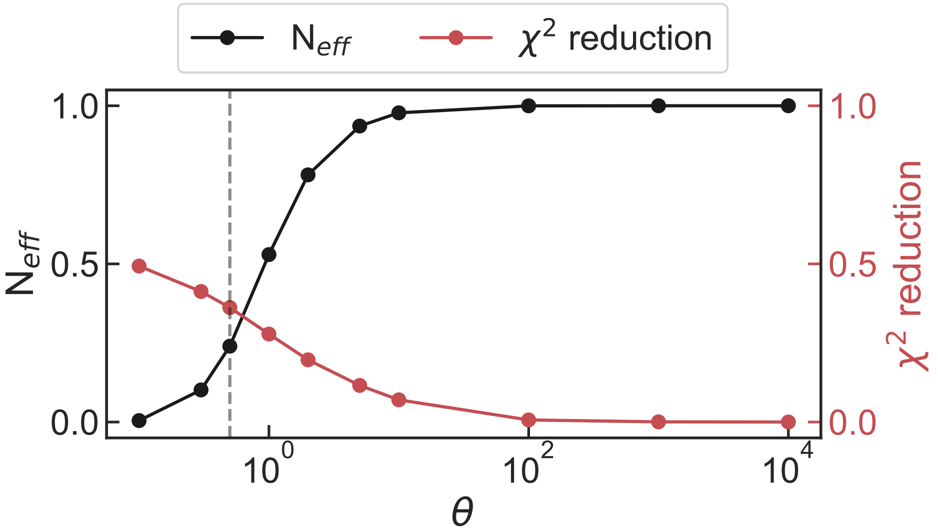


Figure S8: Distribution of the number of effective frames and $\chi^2$ reduction in function of different values of $\theta$. Values obtained for the 1ZE9 model. The selected value of $\theta$ is shown by the vertical line.

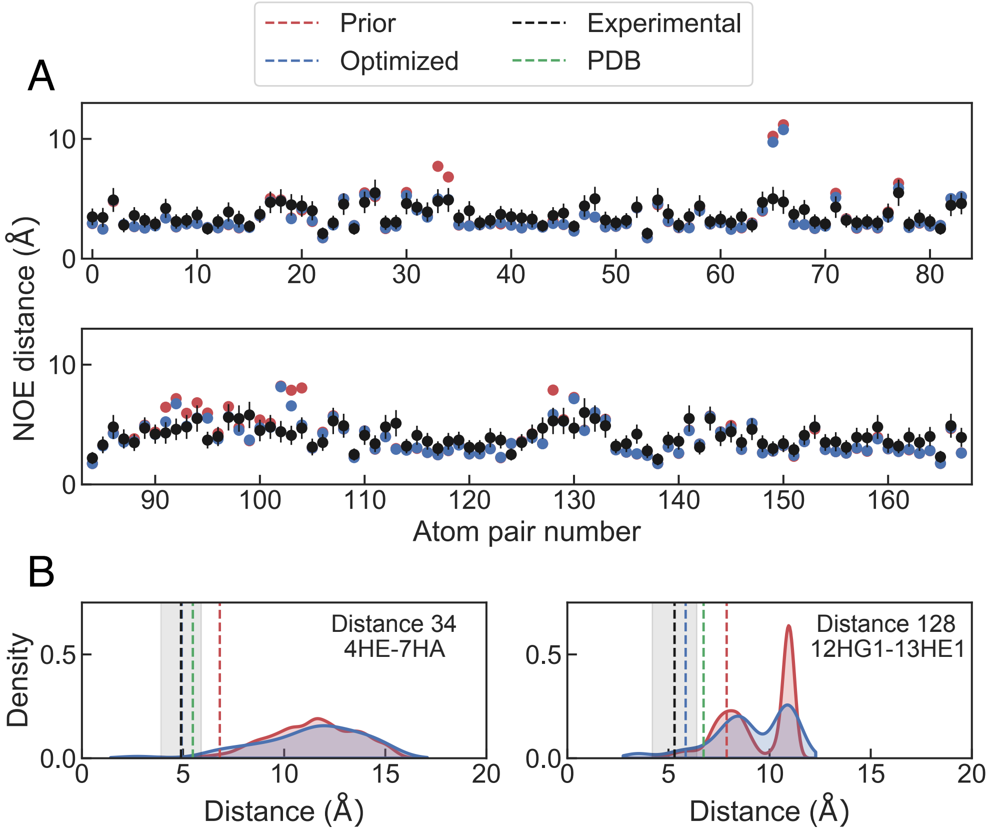


Figure S9: Experimental NOE distances and values back-calculated from simulations (A) and distance distributions for two optimized distances (B) of system PDB ID 1ZE9. Experimental values are shown in black, experimental uncertainty is shown in grey highlight, values extracted from the unoptimized simulation are shown in red, values extracted from the optimized simulation are shown in blue and values calculated from the PDB structures are shown in green.

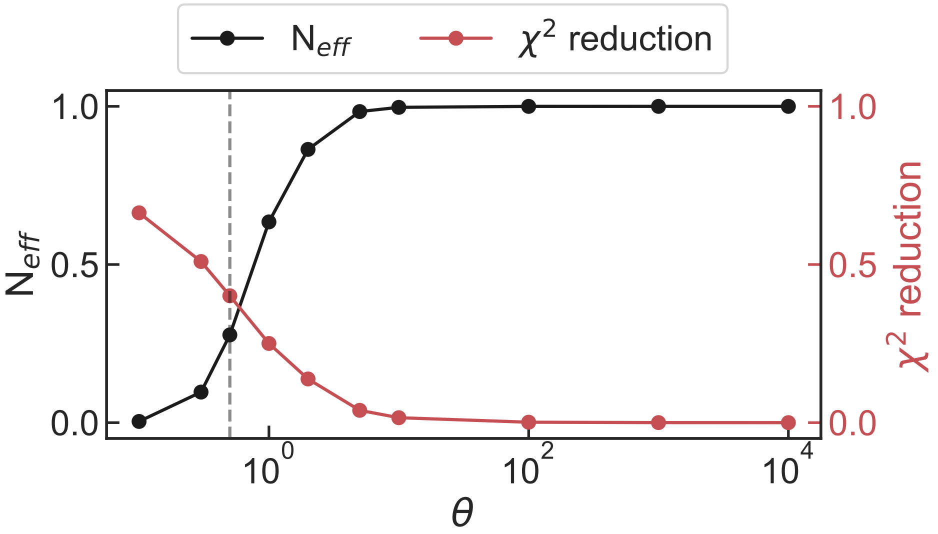


Figure S10: Distribution of the number of effective frames and $\chi^2$ reduction in function of different values of $\theta$. Values obtained for the 5LFY model. The selected value of $\theta$ is shown by the vertical line.

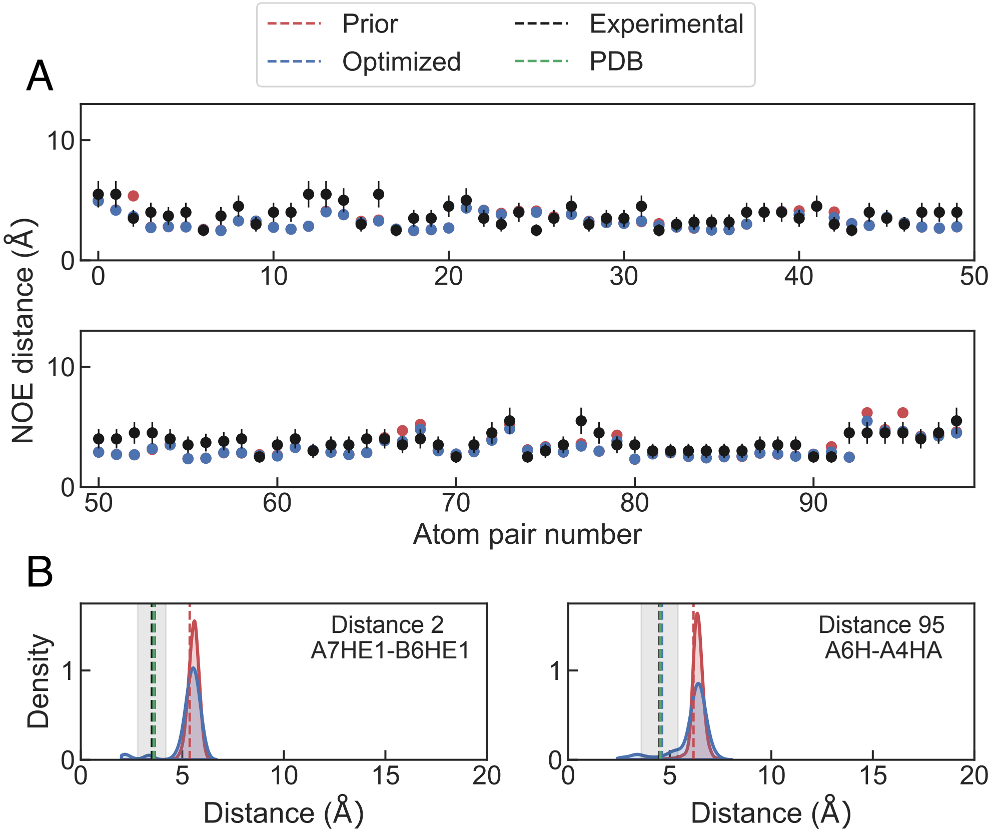


Figure S11: Experimental NOE distances and values back-calculated from simulations (A) and distance distributions for two optimized distances (B) of system PDB ID 5LFY. Experimental values are shown in black, experimental uncertainty is shown in grey highlight, values extracted from the unoptimized simulation are shown in red, values extracted from the optimized simulation are shown in blue and values calculated from the PDB structures are shown in green.

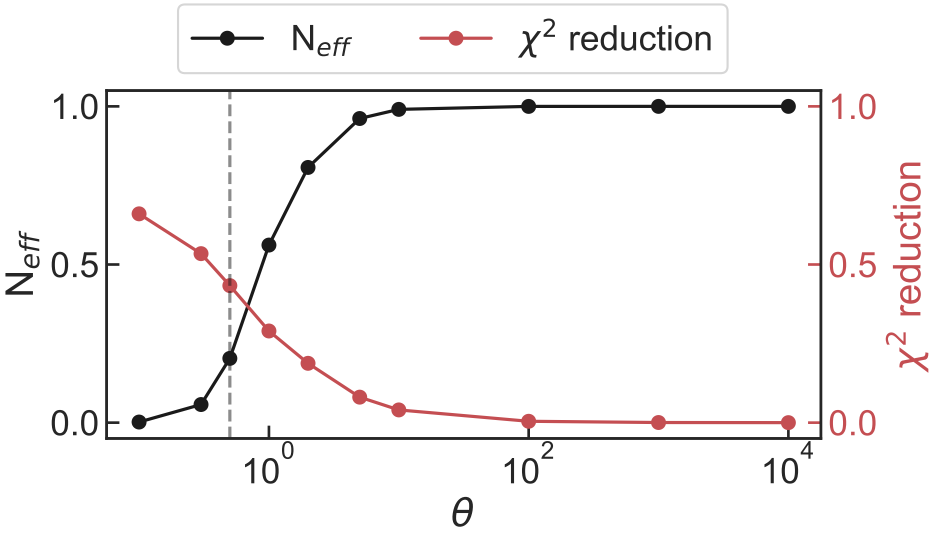


Figure S12: Distribution of the number of effective frames and $\chi^2$ reduction in function of different values of $\theta$. Values obtained for the 2LI9 model. The selected value of $\theta$ is shown by the vertical line.

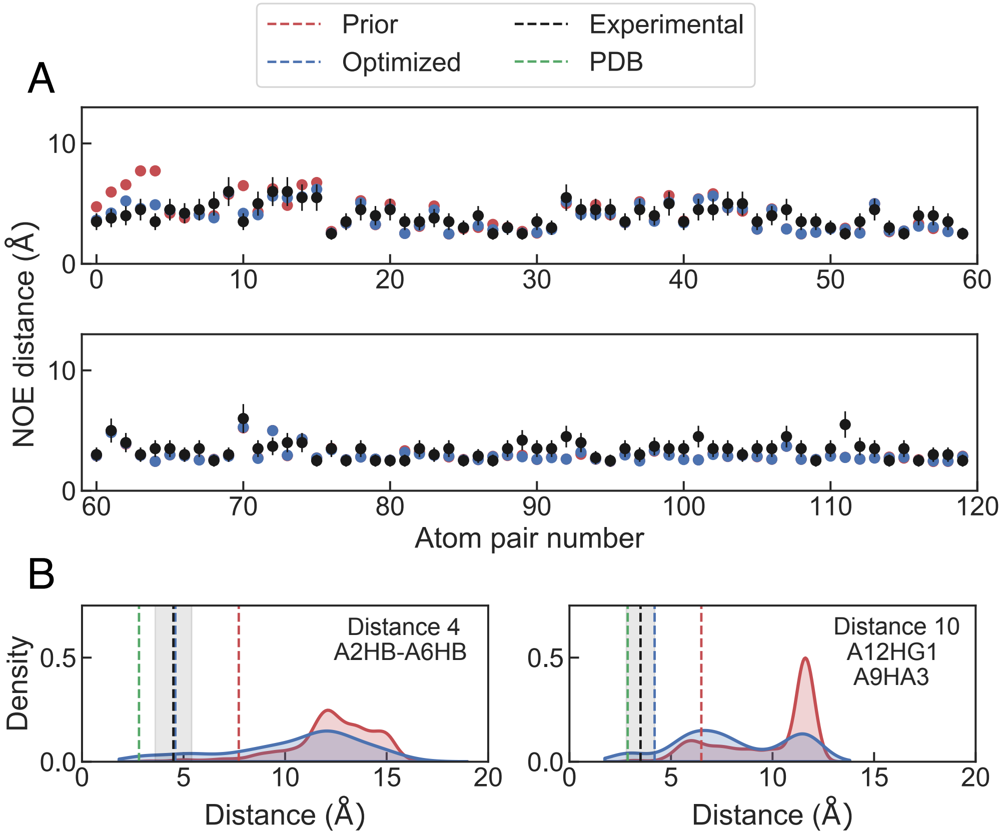


Figure S13: Experimental NOE distances and values back-calculated from simulations (A) and distance distributions for two optimized distances (B) of system PDB ID 2LI9. Experimental values are shown in black, experimental uncertainty is shown in grey highlight, values extracted from the unoptimized simulation are shown in red, values extracted from the optimized simulation are shown in blue and values calculated from the PDB structures are shown in green.

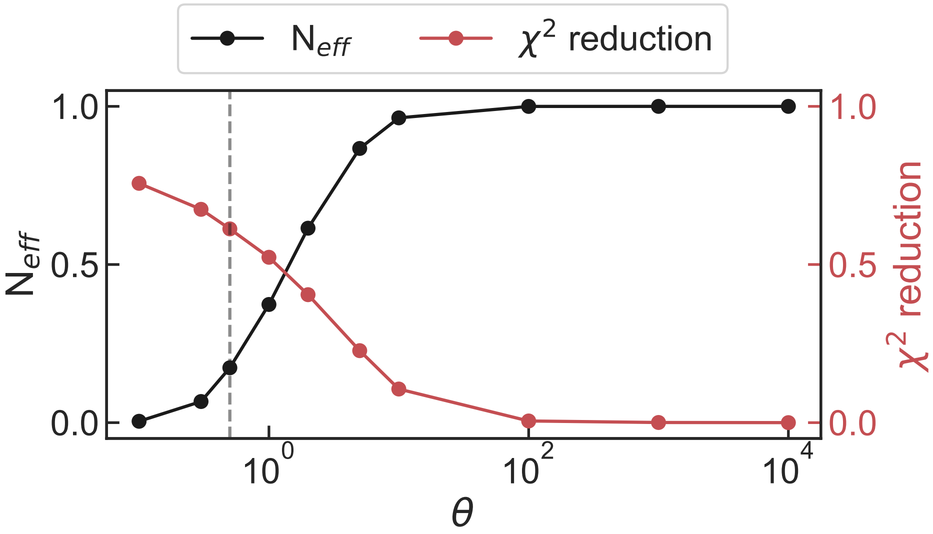


Figure S14: Distribution of the number of effective frames and $\chi^2$ reduction in function of different values of $\theta$. Values obtained for the 2MGT model. The selected value of $\theta$ is shown by the vertical line.

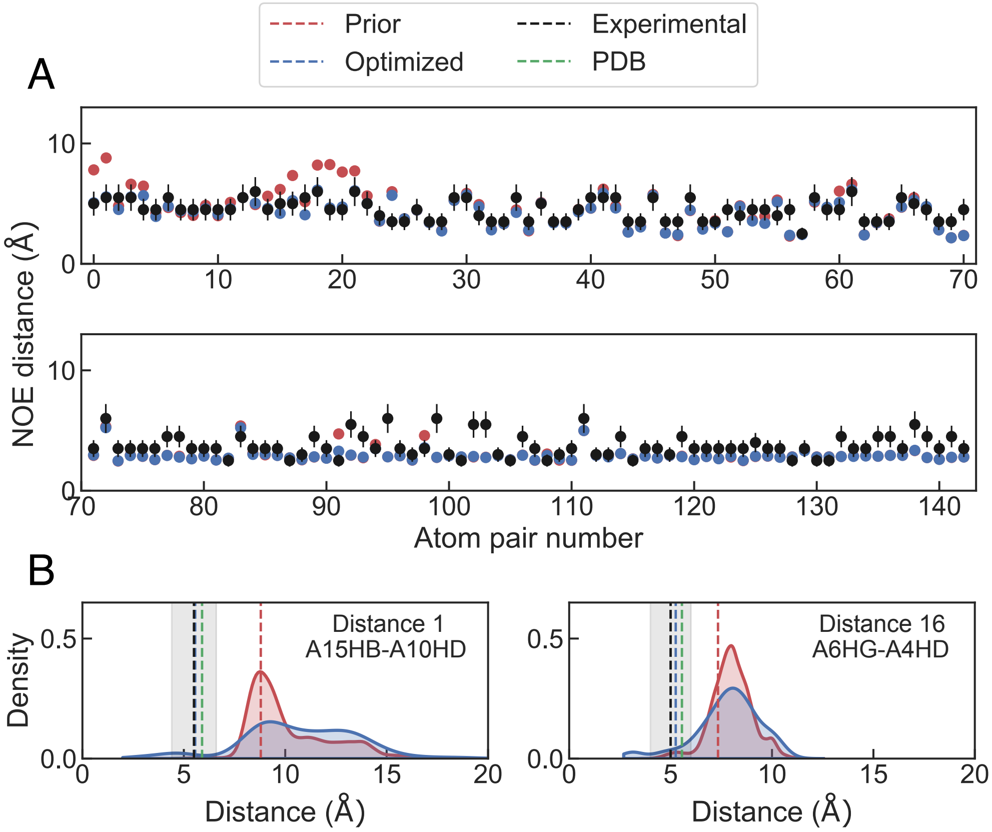


Figure S15: Experimental NOE distances and values back-calculated from simulations (A) and distance distributions for two optimized distances (B) of system PDB ID 2MGT. Experimental values are shown in black, experimental uncertainty is shown in grey highlight, values extracted from the unoptimized simulation are shown in red, values extracted from the optimized simulation are shown in blue and values calculated from the PDB structures are shown in green.

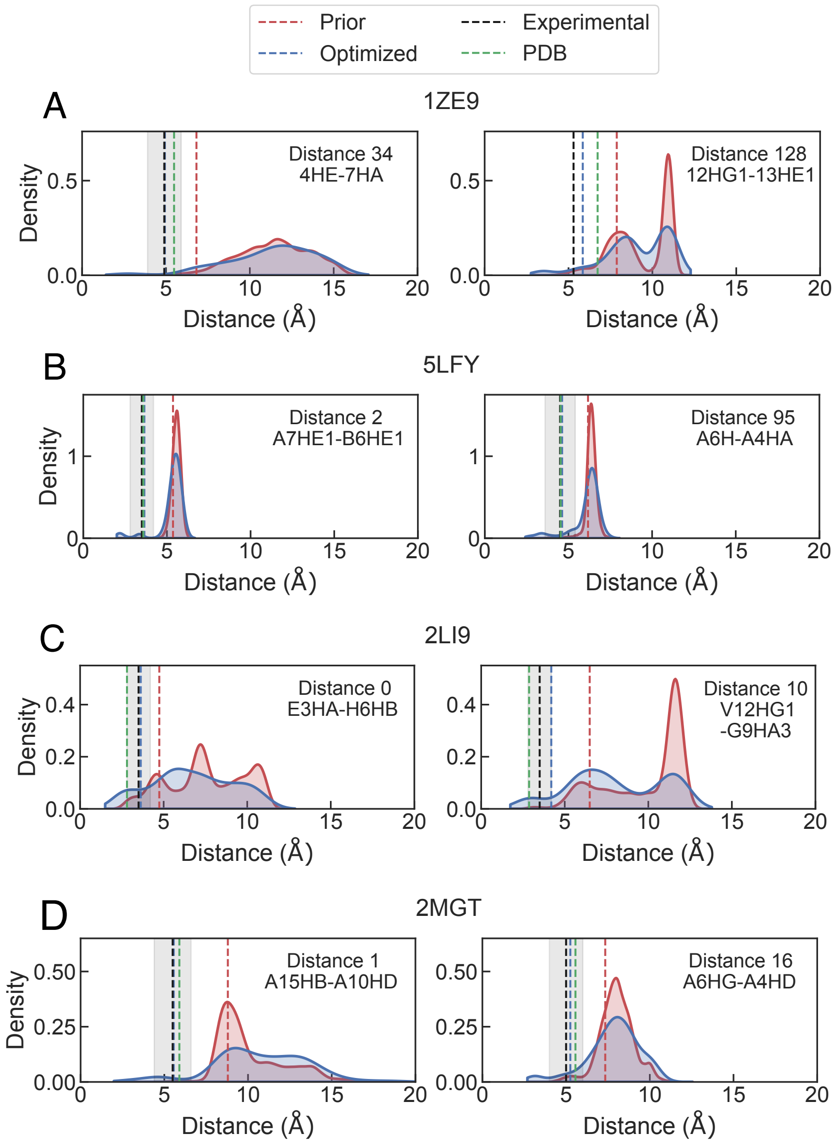


Figure S16: Distribution of distance density of two optimized NOE distances for systems 1ZE9 (A), 5LFY (B), 2LI9 (C) and 2MGT (D). Experimental values are shown in black, experimental uncertainty is shown in grey, values extracted from simulations are shown in red, and reweighted values are shown in blue. The values calculated from the PDB structures are shown in green.
